# Mature larvae continue calling at night in *Vespa mandarinia* from laboratory observations

**DOI:** 10.1101/2024.09.01.610616

**Authors:** Haruna Fujioka, Tatsuya Saga

## Abstract

Animals produce and perceive vibroacoustic signals. *Vespa* hornet larvae produce a rhythmic ‘rasping’ sound by rubbing their mandibles against the cell wall of the nest. The call is thought to be a larval provisioning cue. However, detailed observations of larval calls have been limited to a few species, and it is not known whether the call can be influenced by the external environment, such as light and time of day, or by internal larval states, such as feeding. We conducted laboratory observations of larval calls under workerless conditions to investigate the effects of 1) larval stage, 2) daily variation, 3) light conditions, and 4) feeding treatment. *Vespa mandarinia* larvae produce sounds regardless of their situation, such as position, light condition, time of day, and worker absence. During mastication, the larvae stopped calling. A key finding of this research is the novel discovery that larvae produce sounds at night, which is a previously undocumented behaviour.

## INTRODUCTION

Animals produce and perceive vibroacoustic signals (Greenfield 2016; Virant-Doberlet et al. 2023). Insects, including more than 200,000 species, utilize vibrational communication through various means, such as body and wing movements, high-frequency muscle contractions, and scraping mandibles (Cocroft and Rodríguez 2005; Virant-Doberlet et al. 2023). In social insects, vibrational signals play an important role in communication among nestmates. They have many types and functions in vibrational communication (Hunt and Richard 2013), such as mating (Conrad and Ayasse 2019), foraging (Hrncir and Barth 2014), alarm signalling (Hager and Kirchner 2013; Bota et al. 2022), adult–larva communication (Pepiciello et al. 2018) and larval development (Suryanarayanan et al. 2011).

Vespidae, such as hornets (Vespinae) and paper wasps (Polistinae), make nests from soft paper pulp with sturdy hexagonal cells. The material and shape are favourable for the transmission of vibrational signals (Hunt and Richard 2013). *Polistes* wasps exhibit three types of body oscillation behaviours: antennal drumming, abdominal wagging, and lateral vibration. These types of signals are used for communication among adults and from adults to larvae (Pratte and Jeanne 1984; Suryanarayanan et al. 2011). In *Vespa* hornets, not only adults but also larvae produce sounds via their mandibles, which is a well-known vibroacoustic signal (Schaudinischky and Ishay 1968; Ishay and Landau 1972; Ishay and Schwartz 1973; Ishay et al. 1974; Ishay 1975; Ishay and Brown 1975; Yamane 1976; Edwards and Others 1980; Matsuura and Yamane 1990). *Vespa* larvae rub their mandibles against their cell walls and produce a rhythmic “rasping” sound. Rasping sounds have been reported mainly in *Vespa crabro* (Matsuura 1974), *Vespa orientalis* (Ishay and Landau 1972; Ishay 1975), and *Dolichovespula sylvestris* (Weyrauch 1935; Jeanne 1980). Yamane (1976) reported that “this behaviour has been confirmed in almost all the *Vespa* species observed by me”. *D. sylvestris* and *Vespula* wasps also produce scraping sounds, but not all *Vespula* species do (Weyrauch 1935; Jeanne 1980; Matsuura and Yamane 1990). Although male, queen, and worker larvae produce this sound, fourth- and fifth-instar larvae of *V. orientalis* produce this sound (Ishay and Landau 1972).

The function of sound is known to be a “hunger” signal to workers (Schaudinischky and Ishay 1968; Matsuura and Yamane 1990). In *V. orientalis*, when workers receive recorded vibration sounds, they exhibit brood-care-like behaviour (Schaudinischky and Ishay 1968). *Vespa* workers forage during the daytime (Edwards and Others 1980). If there is no new food for the larvae at night, do the larvae stop calling? In addition, a previous study reported that “only after receiving food does the larva stop emitting “hunger” signals, but it resumes them as soon as it has finished masticating” in *V. crabro* (Ishay and Schwartz 1973). However, knowledge of larval calls is very limited to a few species, except that mature larvae produce sounds (Yamane 1976). *V. mandarinia* is the world’s largest hornet, and the species has recently invaded North America (Nuñez-Penichet et al. 2021). While the behaviour of adults—such as their group predation on honeybees—is well documented (Matsuura and Sakagami 1973), the details of larval communication within the nest, such as their calling, have not been thoroughly investigated. Thus, this study aims to confirm the universality of the larval call and to reveal when *V. mandarinia* larvae produce sounds. We described the first detailed observation of this larval behaviour in *V. mandarinia*. To investigate the effects of 1) the larval stage, 2) daily variation, and 3) mastication, we conducted laboratory observations of larval calls in the absence of workers.

## METHOD

### Collection and laboratory setup

*Vespa mandarinia* is a common hornet in East Asia (Archer 1995). They are commonly found from open fields to low mountains across Japan, excluding the area south of Amami Oshima. We collected four colonies (VM1-4) from 2023-10-20 to 2024-10-2 in Toyota, Aichi Prefecture, Japan. The number of subjected larvae and their caste/stage and colony collection date (day 0) are summarized in Table 1. The nest combs were placed upside down inside an incubator and kept at 25°C in the dark during the nonexperimental period. Food was not provided to the larvae until Day 6. The experimental schedule is shown in Supplementary Figure 1. Adult workers were not included in this study for safety reasons. We labelled each larva by position and took note of the larval stage on the basis of head width and colour (under the 4th and 5th stages). The 5th-instar larvae and 4th-instar larvae were distinguished by their head colour (Figure 1). The survival of the larvae was determined by ascertaining whether the larvae moved after they were tapped with tweezers.

**Table 1.** Summary of larvae and experiments.

| Colony ID | Collection date | Number of queen larvae |  | Number of worker larvae |  | Group-level recording | Individual-level recording |  |  |
| --- | --- | --- | --- | --- | --- | --- | --- | --- | --- |
|  |  | Fifth-instar | Under 4th-instar | Fifth-instar | Under 4th-instar |  | Comparison of larval stage | Observation through out day | Feeding treatment |
| VM1 | 2023-10-20 | 17 | 8 | - | - | yes | yes | yes | yes |
| VM2 | 2024-9-5 | - | - | 26 | 21 |  | yes |  |  |
| VM3 | 2024-9-28 | 15 | 0 | - | - | yes |  | yes |  |
| VM4 | 2024-10-2 | 14 | 7 | 11 | 5 | yes | yes | yes | yes |

**Figure 1.**
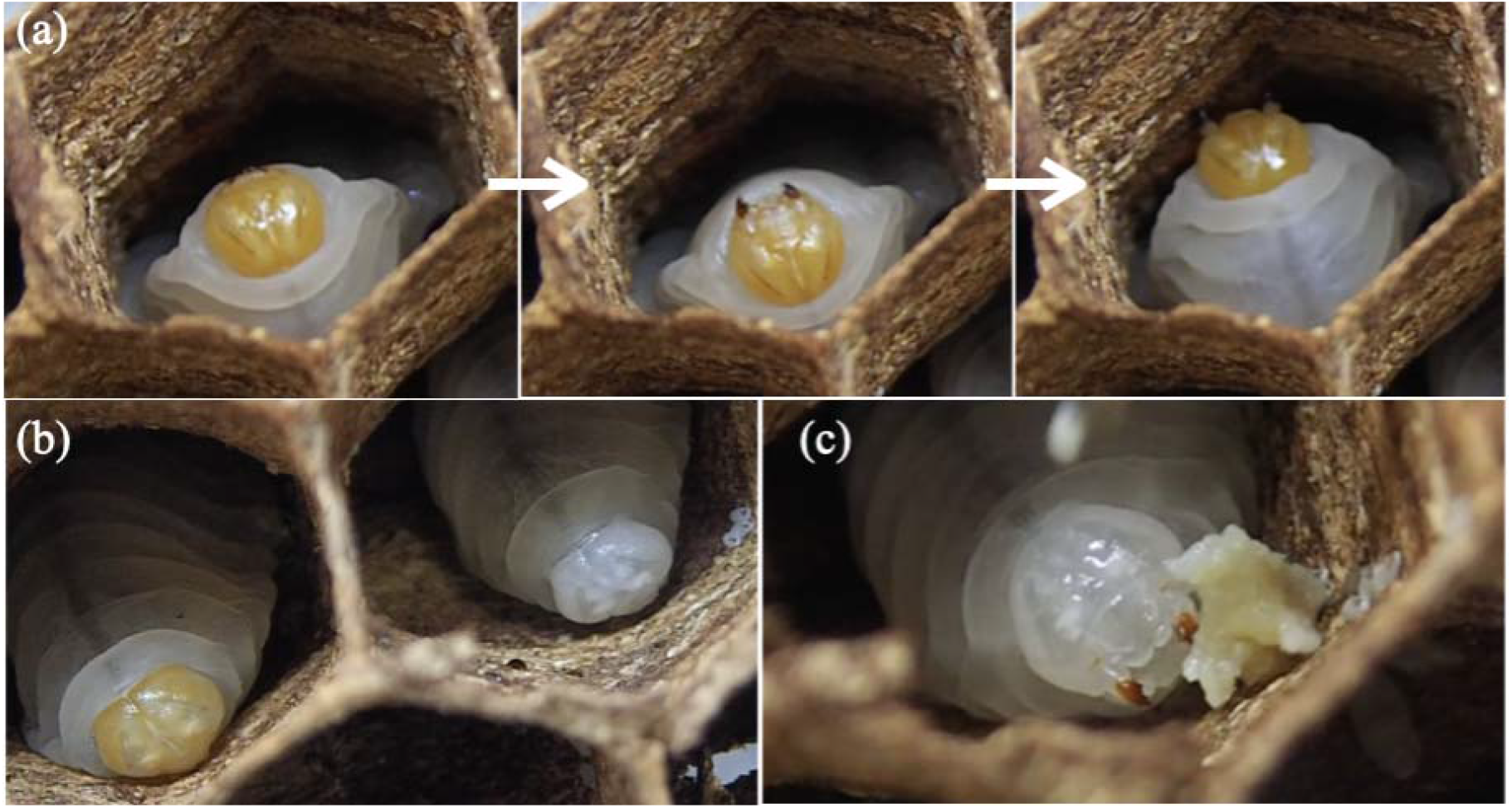
Larval call and stages of *Vespa mandarinia*. Fifth-instar larvae rub their mandibles against their cell walls and produce a rhythmic ‘rasping’ sound, as recorded on October 23, 2023 (a). The number of calls per minute for fourth-instar and fifth-instar larvae. Larvae of *Vespa mandarinia* on October 24, 2023 (b-c). The larva on the left with a yellow head is a fifth-instar larva, and the larva on the right with a white head is a fourth-instar larva (b). A fourth-instar larva eating a piece of cricket (c). Photograph credit: Haruna Fujioka.

### Recording: group level call

We measured any differences in group-level calls between day and night. Additionally, to investigate whether light-on/light-off conditions affect their calls, a group-level recording of calls for 30 minutes was conducted under different light conditions, using three combs that included queen larvae (see Table 1, VM1, VM3 and VM4). The observation dates varied among colonies; for details on the timing of the observations, see Supplementary Figure 1. The nest comb was placed upside down inside an incubator at 25°C. After the group was recorded under dark conditions for 20 minutes, the group recording under light-on (white light) conditions was conducted for 20 minutes. The continuous group-level recordings with different light conditions (dark and light) were conducted twice, between 2 pm and 5 pm [daytime] and between 8 pm and 10 pm [nighttime]. A mic (Yeti, BM400S, logical) and Quick Time Player (version 10.5 on MacBook Pro, OS13.5) were used for the recording. White noise was removed via noise reduction on Audacity (version 3.4.2). WaveSurfer (version 1.8.8p6) was used to plot a spectrogram. After recording, the number of group-level calls was counted every minute for 20 minutes (n = 20) to listen to the sound recorded by an observer.

### Observation: individual-level call

Larval calls were manually counted under light-on conditions. The comb was placed upwards on a table. We used a handy counter and a digital timer to count the total number of calls per minute. Each larva’s calls were counted twice, and the average number of calls was used for data analysis. A set of manual observations was performed within 30 min. To investigate the effect of the larval stage on the number of calls, we compared the number of calls recorded from 12 PM to 6 PM between 5th-instar larvae and those under 4th-instar larvae via VM1, VM2, and VM4. VM3 was not included in the analysis because all larvae under the 4th instar died on the observation start date (Day 5). To investigate the daily variation in the number of calls, we conducted every six hours of observation from 12 pm to 6 pm the following day (six time points), using the VM 1 VM3 and VM4 colonies.

To investigate whether larvae stop calling when they eat, we fed them a piece of cricket and observed the number of calls at three time points on Day 6 via the VM1 and VM4 colonies. After the manual observation of calls per individual at 9 am [before feeding], we fed approximately 0.05 g of defrosted house cricket between 12 pm and 3 pm. We allowed the larvae to feed on crickets ad libitum for 3 hours, although the amount eaten per individual was not controlled. The manual observations of the calls were conducted at two additional times: within 5 minutes after food removal [during eating] and at 3 pm [after the meal].

### Statistics

A t test was used to compare the number of calls between fifth-instar larvae and fourth-instar larvae. A linear mixed model (function *lmer* of package lme4) was used to analyse time of day (daytime or night) and light (light or dark) effects on the frequency of calls per second at the group level. The colony ID was used as a random effect. The differences in the number of calls with respect to daily variation and feeding treatment were compared via the Tukey HSD test.

## RESULTS

We observed that larvae produced a rasping sound when the comb was placed upside down (Figure 1a, Video S1). Even in the absence of workers’ presence around the larvae, the mature larvae continued to call in this study. Larvae continued to make sounds throughout the 30-minute recording regardless of the time of day and light conditions (Figure 2a-d). The number of group-level calls per minute varied among colonies (Figure S2). The number of group-level calls per minute during the daytime was significantly lower than that during the daytime (Figure S2, Table 2, LMM, P < 0.05). The number of group-level calls per minute under the light-on condition was significantly greater than that under the light-off condition (Figure S2, Table 2, LMM, P < 0.05).

**Table 2.** Summary of larvae and experiments.

|  | Estimate | Std. Error | df | t value | P value |
| --- | --- | --- | --- | --- | --- |
| (Intercept) | 0.5570 | 0.058 | 2 | 9.56 | 0.01 |
| Light (light-on) | 0.0268 | 0.0069 | 235 | 3.83 | < 0.001 |
| Time of day (night) | 0.0223 | 0.0069 | 235 | 3.19 | < 0.001 |

**Figure 2.**
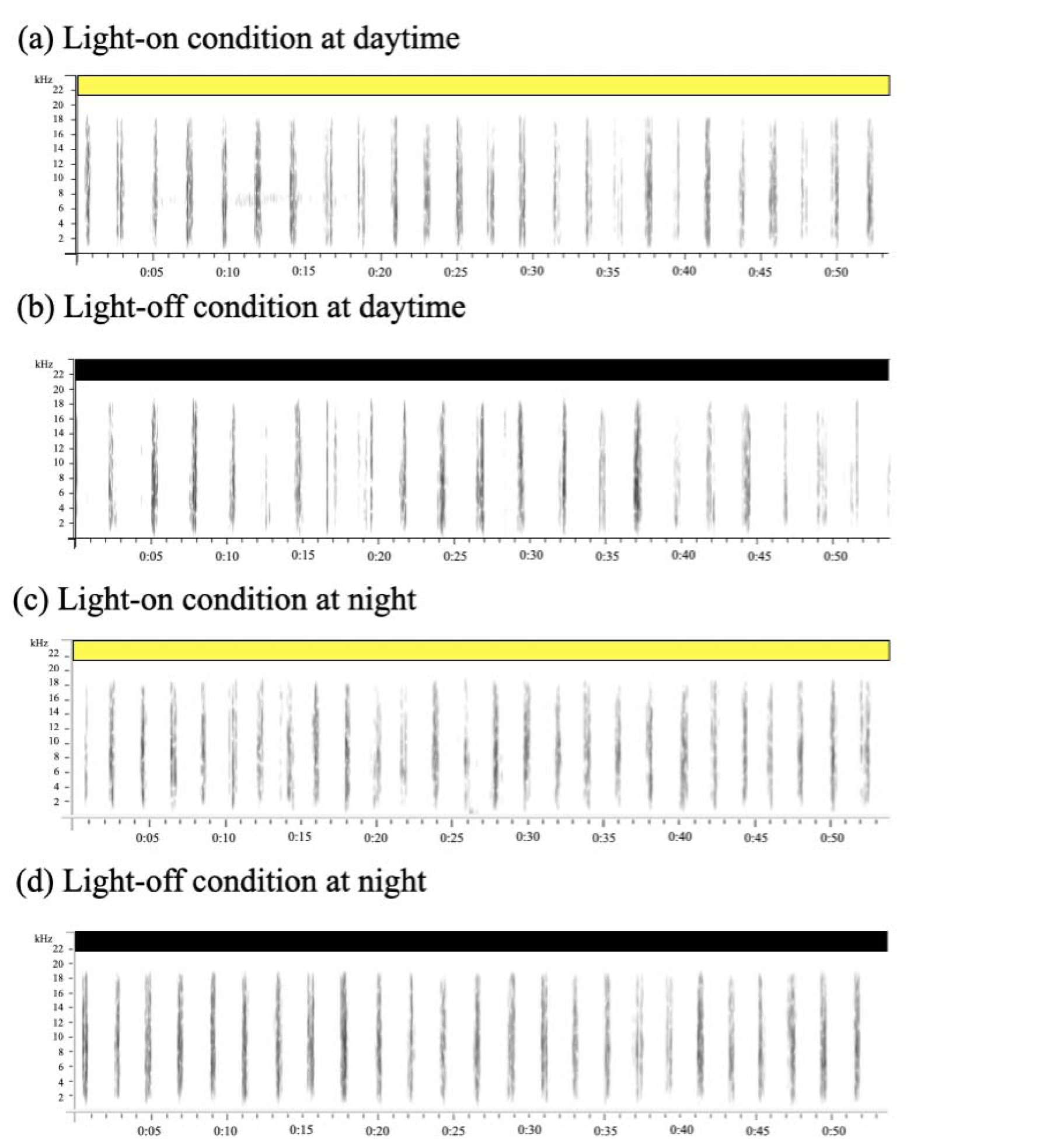
Spectrogram of group-level calls. Spectrograms of group-level calls under light conditions (a, c) and dark conditions (b, d). Daytime observations were made between 2 pm and 5 pm, and nighttime observations were made between 8 pm and 10 pm. The y-axis indicates the frequency (Hz).

Only fifth-instar larvae continuously produced sounds (Figure 3 and Video S1). The number of calls by fifth-instar larvae was significantly greater than that by fourth-instar larvae in all colonies and on all days (Figure 3, Wilcox test, P < 0.05). We confirmed the viability of larvae under fourth instar conditions by observing that they moved when we provided food (Figure 1c) or physical stimuli (tapping them with tweezers).

**Figure 3.**
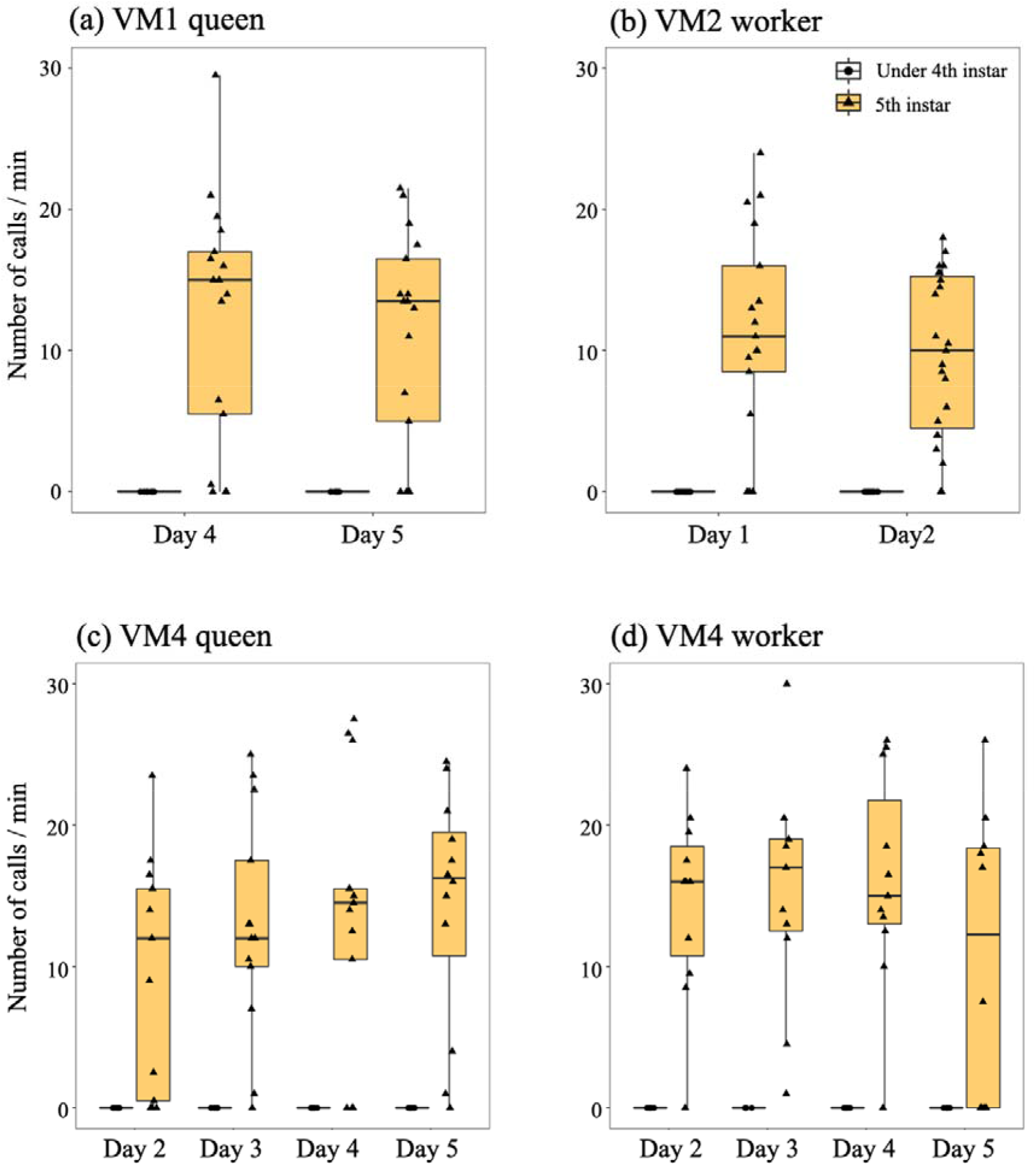
Effects of the larval stage on larval calls at the individual level. The number of calls was observed during the daytime under light conditions from Day 1 to Day 5.

In three colonies (VM1, VM3, and VM4), there were no significant daily differences in individual-level calls (Figure 4, Tukey-HSD test, P > 0.05). Observations revealed variations in the calls of fifth-instar larvae; some fifth-instar larvae presented fewer calls in VM1 and VM3. Additionally, during the five-day observation period, despite the absence of food and water, some 5th instar larvae perished, whereas others survived and continued to emit calls.

**Figure 4.**
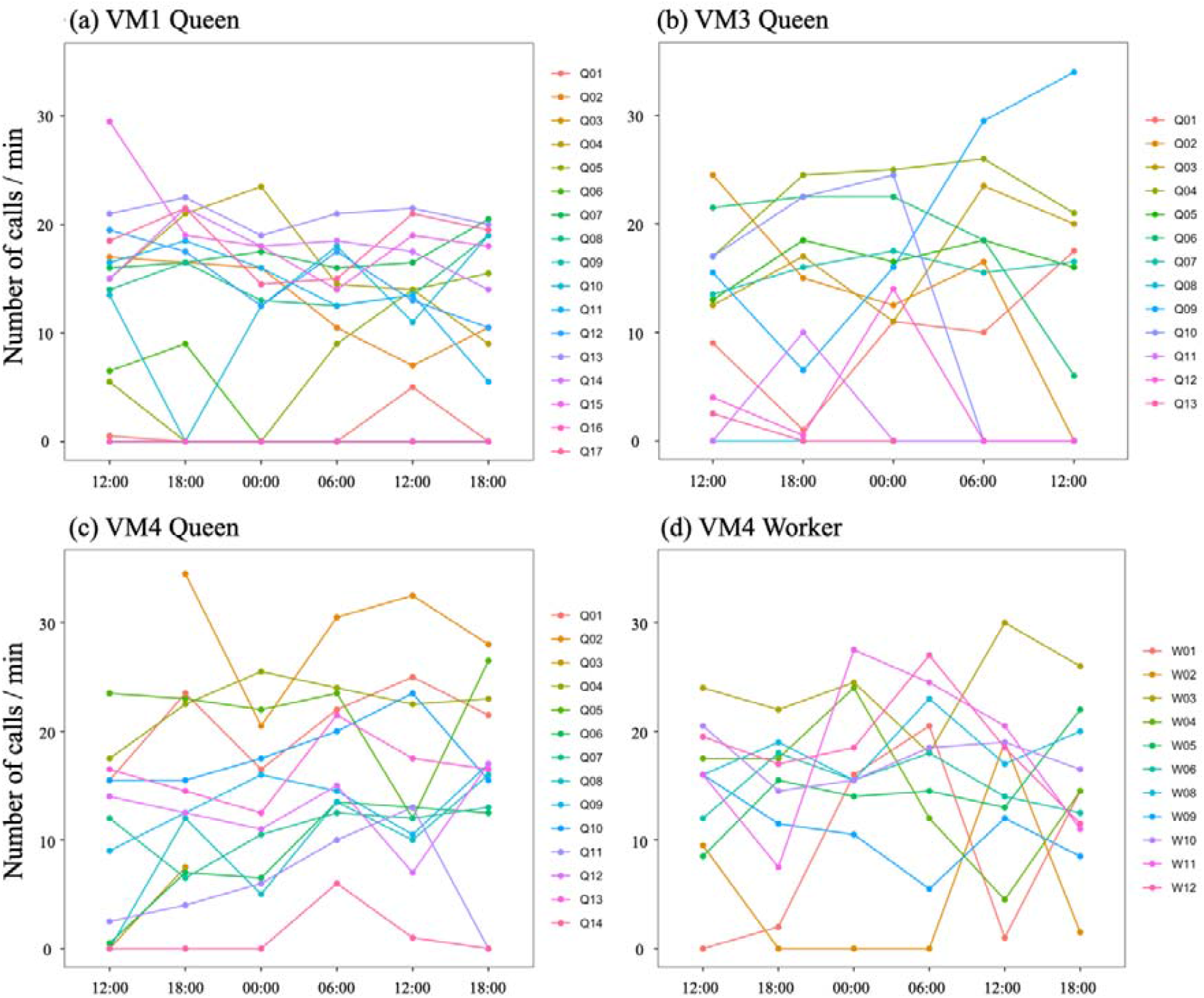
Daily pattern of the number of calls at the individual level. The number of calls at the individual level was counted every 6 hours from 12 am. Each point represents an observation. The colour indicates larval identity.

The number of calls did not differ before and after feeding (Figure S3, Tukey HSD test, P > 0.05). There was large individual variation in the number of calls immediately after food removal in VM 1 (Figure S3). During mastication, many larvae stopped calling in VM4 (Figure S4, Tukey HSD test, P < 0.05). After food removal, the larvae turned their bodies around, rapidly opened and closed their mandibles, and resumed calling.

## DISCUSSION

This study provides the first detailed observation of larval calls in *Vespa mandarinia. V. mandarinia* larvae produced sounds regardless of their situation, such as position, time of day, light conditions, and worker absence. We confirmed that *V. mandarinia* larvae continue calling even when the nest is turned upside down, which is inconsistent with the findings of a previous study of *V. orientalis* (Schaudinischky and Ishay 1968). Sound production depends on the larval stage (Figure 3), which is consistent with previous studies (Ishay and Landau 1972; Yamane 1976; Edwards and Others 1980; Matsuura and Yamane 1990). Our results revealed that in the 4th early stage, larvae did not produce sounds (Figure 3). For these early-stage larvae, prioritizing energy conservation for growth could be crucial. In addition, some fifth-instar larvae do not produce sounds (Figure 3). There may be three possible factors. First, larvae have soft mandibles immediately after moulting, which may prevent them from producing sound. Second, if starvation is severe, larvae may become immobilized. After observation on Day 6, some larvae died while turning black (Figure S5). Third, fifth-instar larvae, which have reached sufficient maturity and readiness to pupate, cease to receive nourishment from workers. This cessation of feeding may result in a corresponding cessation of sound production in response to altered conditions. To reveal why some larvae do not perform calling, future observations are needed on the amount of food received and the brood care workers provide to young larvae.

A key finding of this research is that larvae make sounds during the night (Figure 2, 4). The sound production of larvae at night was also observed when some emerged workers presented themselves in their colony (T. Saga, personal observation). *V. mandarinia* is known to be a diurnal hornet; only one species, *V. crabro*, forage after sunset in Japan (Matsuura and Yamane 1990). It is not reasonable that larvae continue “begging” calling at night because there is no new food for larvae at night. In a future study, the time available for workers to feed larvae should be investigated to determine why larvae beg for food at night. One possibility is that these larval calls at night will likely serve an information-sharing purpose for foragers, informing workers about the current number of larvae and food requirements of the following day. Conversely, the increase in call frequency with artificial illumination could be influenced by altered environmental conditions, deviating from the natural state of darkness. Further detailed observations under conditions closer to natural conditions could validate these possibilities, enhancing our understanding of larval communication dynamics.

We found that during mastication, larvae stopped calling (Figure S3, S4). After the food was removed, the larvae resumed sound. One of the main problems of this study is that we did not sense actual larval hunger levels. It is unknown how much individual larvae eat and what nutrients they need. Vespine wasps use various food resources, both liquid foods and solid flesh pellets (Matsuura and Yamane 1990). Their diet consists of large insects such as longhorn beetles, scarab beetles, mantises, caterpillars of Lepidoptera, and large orb-weaving spiders <u>(Matsuura 1984, 1991</u>, Matsuura and Yamane 1990). The food given was only a small piece of cricket, which is a protein-rich food. The protein-biased food and three hours of feeding may not have been enough to ensure that the larvae were full after 5 days of starvation. Therefore, these data do not suggest that starvation levels influence the production of calls. Additional experiments controlling the amount of food are necessary.

We observed that fifth-instar larvae can survive without food or water for up to five days, not only persisting but also continuing to produce rasping sounds. Hornet larvae not only receive food from adults but also contribute to the nest’s food storage function by regurgitating the food back to the adults (Maschwitz 1966), an aspect that highlights their resilience to starvation. As these larvae are in the prepupal stage, there is variation in body size among individuals, suggesting that larger individuals may have greater resistance to starvation. Identifying the correlation between body size and starvation resistance remains a subject for future research. An interesting new question related to body size is the differences in the frequency and strength of calls among queens, males and workers. The body size of a new queen is a critical determinant of successful overwintering and colony foundation (Saga et al. 2024). Specifically, in the realm of independently founding social insects, more prominent queens possess a competitive advantage in founding new nests (Keller 1994; Heinze and Tsuji 1995; Wiernasz and Cole 2003). Reproductive individuals may produce stronger and greater numbers of calls than workers do.

In summary, during more than 24 hours of starvation, fifth-instar *V. mandarinia* continuously called. The novel finding of this research is that larvae produce sounds at night, a previously undocumented behaviour. When larvae eat, they stop making calls, which is consistent with the findings of a previous study (Ishay and Schwartz 1973). Although the production of provisioning cues can be costly, the only thing that larvae can do in the nest is beg for food.

## ACKNOWLEDGMENTS

We would like to thank Fumihiro Sato, Tsuneo Toyama, and Nagashi Fukatsu for their help in collecting the *Vespa mandarinia* used in this study. We are also thankful for research promotion support from the Kobe University Graduate School of Human Development and Environment, TAKEO Corporation, and Ryobi-Teien research grants. We thank Koki R. Katsuhara for his advice about the data analysis.

