## Supplementary materials for "Mature larvae continue calling at night in *Vespa mandarinia* from laboratory observations"

**Supplemental materials**

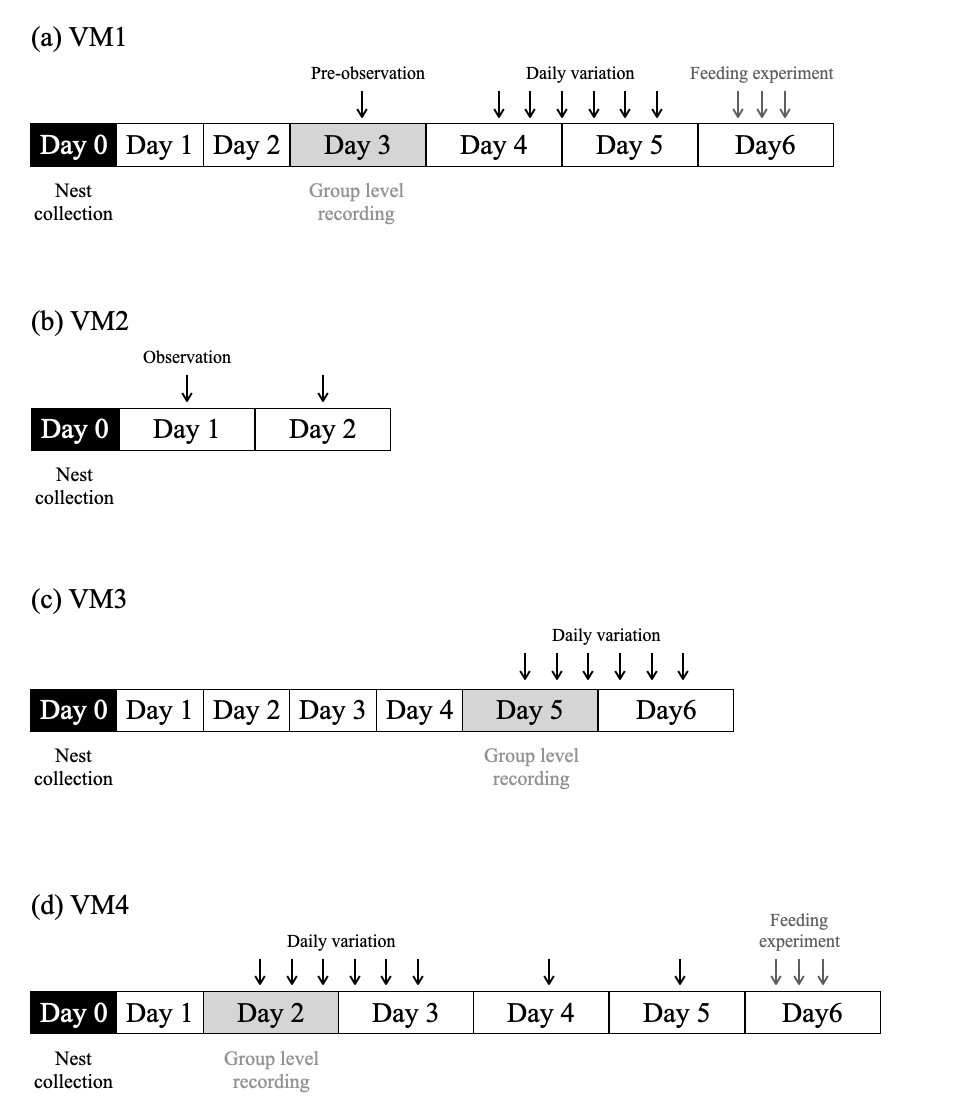
Figure S1. Experimental schedule.

The black arrows indicate the observation of counting the number of larval calls per individual. Day 0 indicates a nest collection date. Group-level recordings were conducted on Day 3 of VM1, Day 5 of VM3, and Day 2 of VM4. The grey arrows indicate feeding experiment.

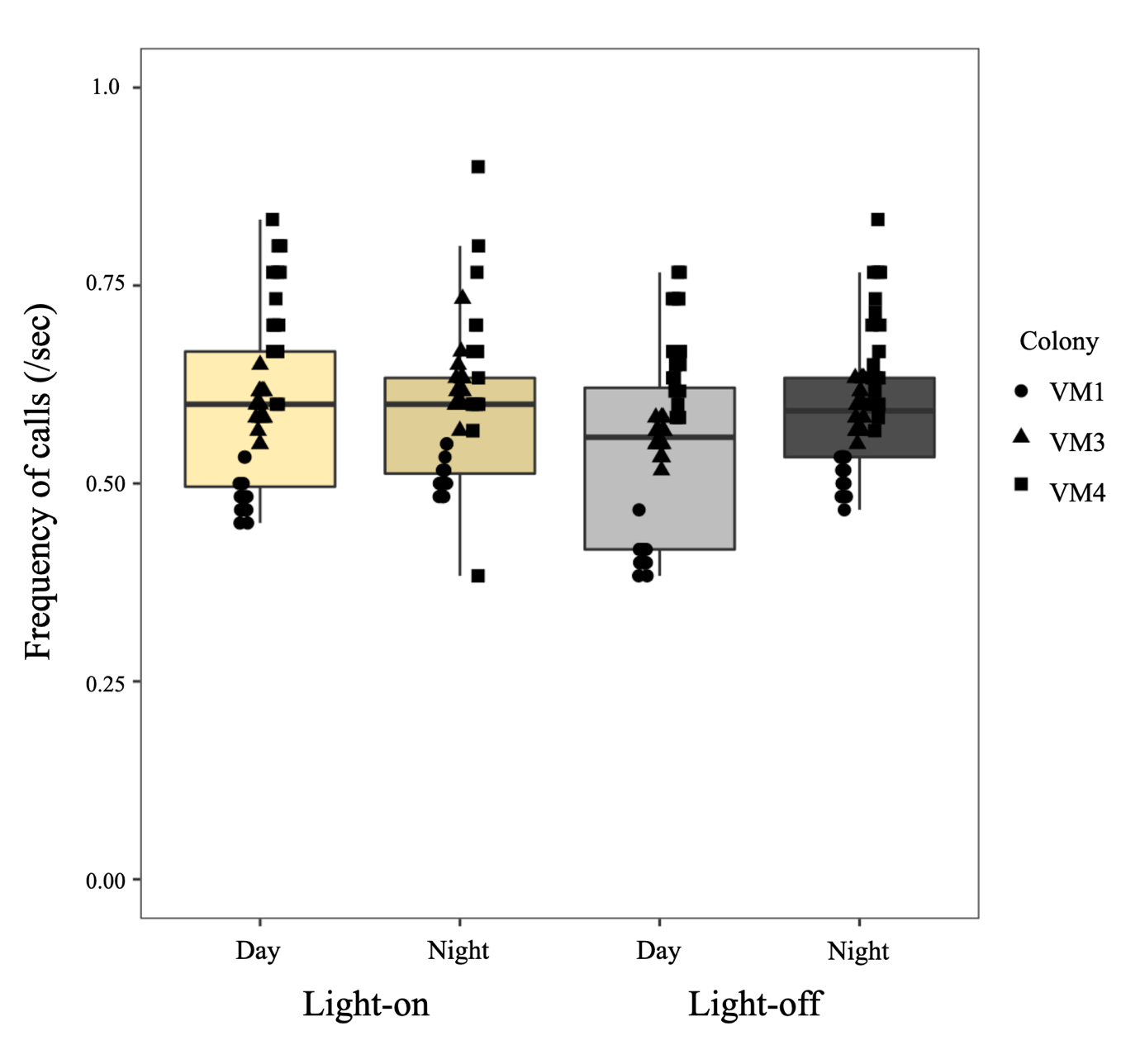

Figure S2. Number of group-level calls of queen larvae per minute under dark and light conditions on Day 3.

The daytime observations were made between 2 pm and 5 pm, and the nighttime observations were made between 8 pm and 10 pm. The yellow and grey boxes indicate light and dark conditions, respectively. Circle, triangle and square indicate different colonies; VM1, VM3, and VM4, respectively

.

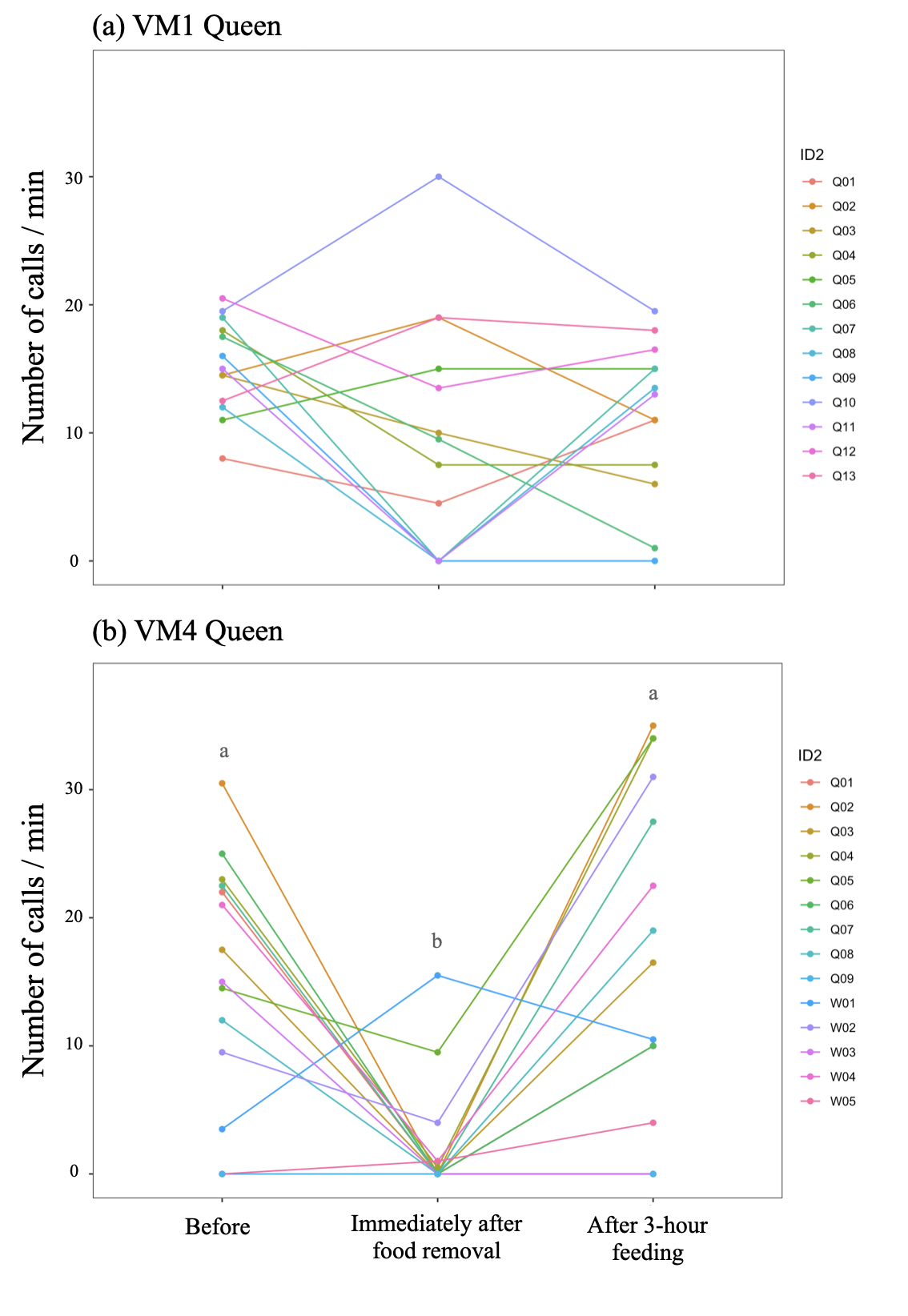

Figure S3. Larvae stopped calling during mastication. The number of calls at the individual level before feeding at 9 am, immediately after removing the food, and after 3 hours of feeding at 3 pm on Day 6. Q and W indicates queen and worker larvae, respectively (b). Different letters on the box indicate significant differences by Tukey-HSD test (a, b: P < 0.05).

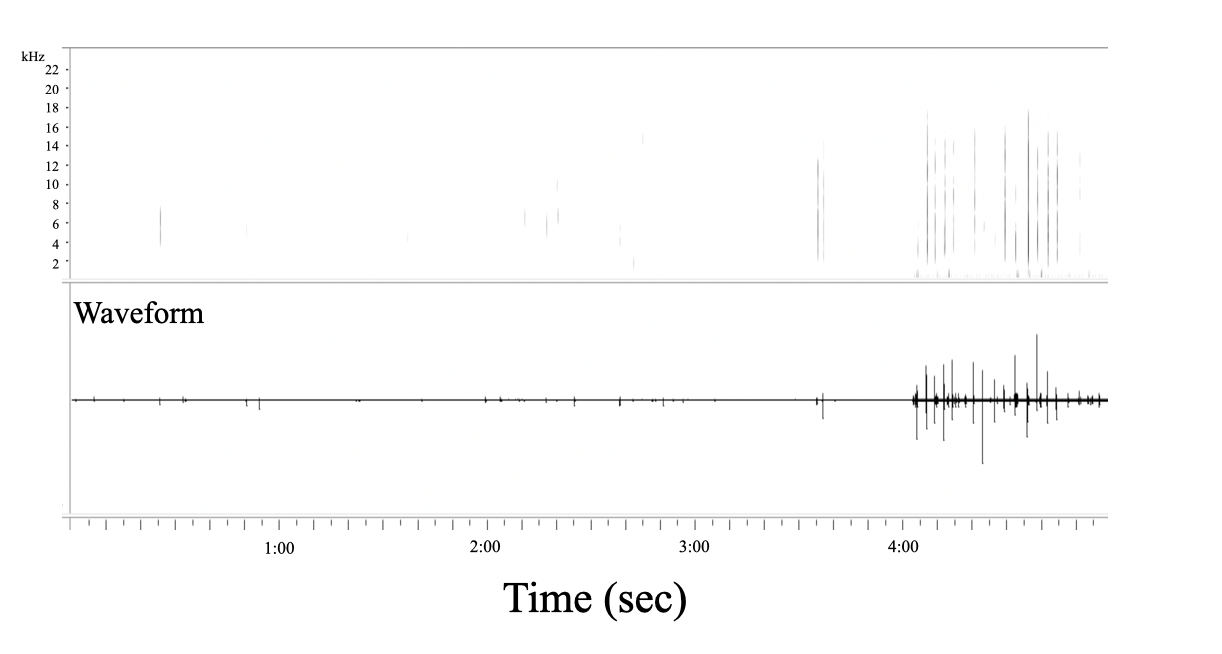

Figure S4. Spectrogram and waveform of group-level calls in the feeding experiment. Larvae stopped calling during mastication.

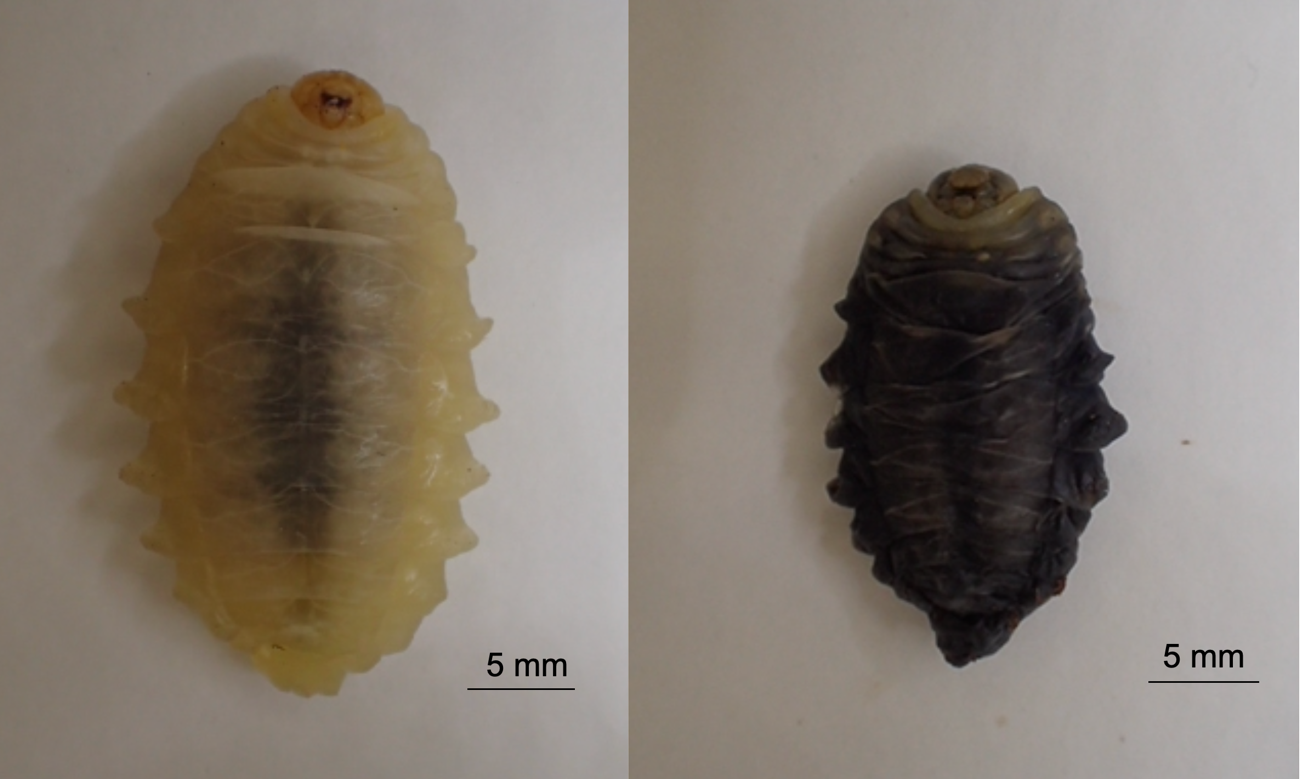

Figure S5. Alive 5-th instar larva (left) and dead 5th-instar larva (right).
